# Endocardial differentiation in zebrafish occurs during early somitogenesis and is dependent on BMP and *etv2* signalling

**DOI:** 10.1101/654525

**Authors:** Samuel J Capon, Kelly A Smith

**Affiliations:** Division of Genomics of Development and Disease, Institute for Molecular Bioscience, The University of Queensland, Brisbane, Queensland 4072, Australia.

## Abstract

The endocardium and adjacent vascular endothelial network share a number of molecular markers however there are distinct physiological functions of these tissues. What distinguishes these lineages on a molecular level remains an important, unanswered question in cardiovascular biology. We have identified the *Gt(SAGFF27C); Tg(4xUAS:egfp)* line as a marker of early endocardial development and used this line to examine endocardial differentiation. Our results show that the endocardium emerges from the anterior lateral plate mesoderm at the 8-somite stage (13 hpf). Analysis in a number of loss-of-function models showed that whilst *nkx2.5*, *hand2* and *tal1* loss-of-function have no effect on the endocardial progenitor domain, both *etv2* loss-of-function and inhibition of BMP signalling reduce the endocardial domain. Furthermore, manipulating BMP signalling alters *etv2* expression. Together, these results describe the onset of endocardial molecular identity and suggest a signalling cascade whereby BMP signalling acts upstream of *etv2* to direct differentiation of endocardial progenitors.

## Introduction

During early stages of embryonic development the heart is comprised of two major tissues: the myocardium and endocardium. To date, much of the research concerning heart development has focused on the myocardium, the beating muscle of the heart, while the endocardium has received relatively little attention. The endocardium is a specialised subset of endothelium that forms the inner lining of the heart. It plays important roles in the development of the trabecular myocardium, contributes to the coronary vasculature, septa and cardiac cushions, and connects the heart with the adjacent vascular endothelial network (Markwald et al. 1975; Markwald et al. 1977; Meyer & Birchmeier 1995; Kisanuki et al. 2001; Stankunas et al. 2008; Harris & Black 2010; Red-Horse et al. 2010; X. Tian et al. 2017). Despite these important and unique roles, the developmental cues that distinguish the endocardium from the vascular endothelium remain to be determined.

Genetically, the earliest specific marker of endocardium is *Nfatc1* (de la Pompa et al. 1998; Wong et al. 2012). The endocardial-specific expression of this marker, combined with the functionally distinct roles of the endocardium, demonstrate that the endocardium is a unique subset of endothelium with its own transcriptional signature. However, when this distinction is first established at a molecular level remains unclear. In mice, *Nfatc1* expression is already detectable in the medial aspect of the cardiac crescent at E7.5 (de la Pompa et al. 1998), whereas in zebrafish, *nfatc1* is reported to initiate at 22 hpf, corresponding with the formation of the linear heart tube (Wong et al. 2012). Prior to heart tube formation in zebrafish, endocardial progenitors bud from bilateral, *kdrl^+^/fli1a^+^*populations at the 12-somite stage (12ss; 15 hpf) and migrate to the midline to form the endocardial core of the cardiac disc (Bussmann et al. 2007). This observation suggests that the endocardial and vascular endothelial progenitors are functionally distinct as early as the 12ss, when this subset of cells begin migrating to the midline. However, molecular evidence that corroborates this functional distinction has not been reported.

One hypothesis to explain this lack of evidence is that the endocardium develops from a general endothelial progenitor population sequestered within the cardiac field. In this model, the endothelial progenitors of the heart become encapsulated by myocardium as the cardiac disc forms. The differentiation of the specialised endocardium from this general endothelial population would then occur through myocardial-to-endocardial signalling. As a result, the molecular distinction between the endocardium and the vascular endothelium would occur at, or after, the formation of the cardiac disc, in agreement with the reported expression of *nfatc1* (Wong et al. 2012). However, this hypothesis doesn’t account for the migration of endocardial, but not endothelial, progenitors to the midline at the 12ss.

The early expression of *Nfatc1* in the mouse supports an alternative hypothesis where the endocardium and the vascular endothelium differentiate as unique lineages from an early developmental stage. In agreement with this model, the zebrafish *cloche* mutant lacks all endocardium, yet retains some endothelium, suggesting that the endocardium and vascular endothelium have distinct developmental origins (Stainier et al. 1995). The *cloche* mutation was recently mapped to the *npas4l* gene (Reischauer et al. 2016). Analysis of *npas4l* expression revealed that expression initiates shortly after gastrulation, at 6 hpf, peaks at the tailbud stage, 10 hpf, and subsequently reduces to undetectable levels by 24 hpf (Reischauer et al. 2016). Together, this evidence supports the hypothesis that endocardial identity is established early in development.

Here, we report the identification and characterisation of a previously reported marker of the zebrafish lymphatic system as a novel marker of early endocardial progenitors (Bussmann et al. 2010; Okuda et al. 2012). Characterisation of this line, and *nfatc1* expression, identified distinct endocardial expression domains as early as the 8ss (13 hpf). These results confirm the hypothesis that the endocardium and vascular endothelium differentiate as unique lineages from an early developmental stage. Furthermore, using a number of loss-of-function models, we identify both *etv2* and BMP signalling as important regulators of endocardial differentiation. Further analysis of these factors suggests that BMP signalling acts upstream of the transcription factor *etv2* to direct differentiation of endocardial progenitors.

## Results

### The *Gt(SAGFF27C); Tg(4xUAS:EGFP)* line is a marker of early endocardial development

We obtained the *Gt(SAGFF27C); Tg(4xUAS:EGFP)* line, originating from a large gene-trap screen (Asakawa et al. 2008), for use as a transgenic reporter of the endothelium. Analysis of 48 hours post fertilisation (hpf) embryos showed fluorescence throughout the endothelium, with stronger fluorescence intensity in the endocardium compared to the adjacent vasculature (figure 1 supplement 1). Examining earlier stages revealed fluorescence in the endocardium at 24 hpf with no obvious signal in the adjacent vascular endothelial network (figure 1 supplement 1), suggesting that, at least at early stages, GFP expression in this line is restricted to the endocardium, providing a novel tool to investigate early endocardial development.

To more carefully characterise the expression of the *Gt(SAGFF27C); Tg(4xUAS:EGFP)* line (henceforth referred to as *Gt(endocard:egfp*)), an immunofluorescence time course was performed on *Gt(endocard:egfp*) embryos crossed to the vascular transgenic line, *Tg(kdrl:Hsa.HRAS-mCherry)* (figure 1). The pan-endothelial *Tg(fli1a:EGFP)* line was also examined as a basis of comparison (figure 1). Expression of *Gt(endocard:egfp*) colocalises with *Tg(kdrl:Hsa.HRAS-mCherry)*, confirming that expression for *Gt(endocard:egfp*) is endothelial (figure 1A). Expression of both *Gt(endocard:egfp*) and *Tg(fli1a:EGFP)* lines was observed in bilateral populations of the anterior lateral plate mesoderm as early as the 10ss (figure 1). The *Gt(endocard:egfp*) line showed restricted fluorescence in these bilateral populations (figure 1A), whereas the *Tg(fli1a:EGFP)* line had a broader expression domain, extending in the anterior of the embryo, to include presumptive vascular endothelium (figure 1B).

**Figure 1.**
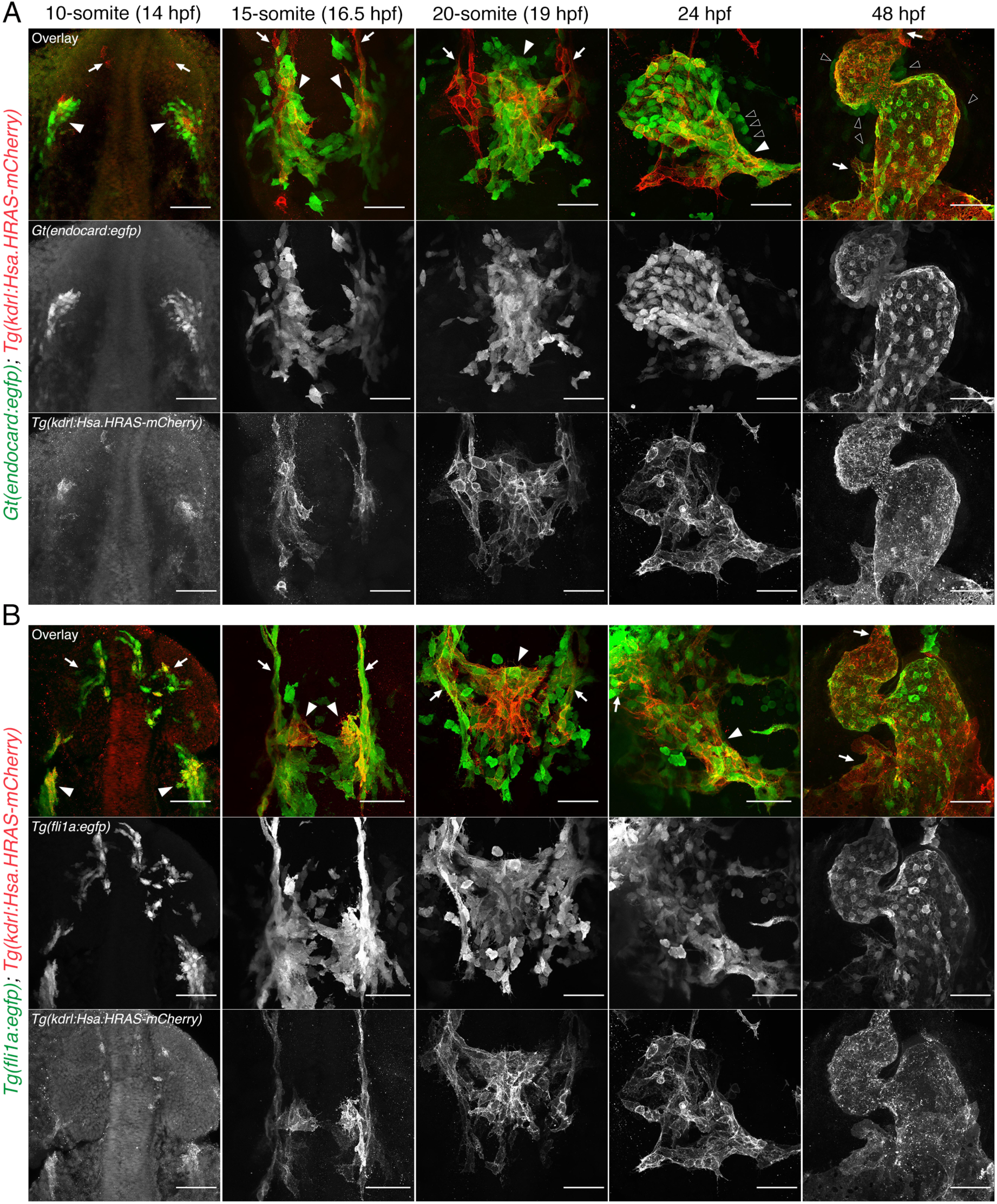
Timecourse analysis of *Gt(endocard:egfp*) shows restricted endocardial expression at somitogenesis stages in the zebrafish embryo. **A.** Immunofluorescence staining of *Gt(endocard:egfp*); *Tg(kdrl:Hsa.HRAS-mCherry)* and **B.** *Tg(fli1a:egfp)*; *Tg(kdrl:Hsa.HRAS-mCherry)* embryos from the 10ss through to 48 hpf. Anterior to the top. Anterior views at 48 hpf, all other images show dorsal views. White arrowheads label fli1a^+^/kdrl^+^/SAGFF27C^+^ endocardial cells. White arrows label fli1a^+^/kdrl^+^/SAGFF27C^-^ vascular endothelial cells. Black arrowheads label presumptive myocardial cells. Scale bars represent 50 µm.

Consistent with previous reports (Bussmann et al. 2007), the bilateral populations of endothelium were found to have migrated to the midline by the 15ss, (figure 1) and by 20-somites have fused to form the endocardial core of the cardiac disc (figure 1). In both crosses, a number of GFP^+^/mCherry^-^ cells are observed around the cardiac disc. Live timelapse imaging of the *Gt(endocard:egfp*) and *Tg(fli1a:EGFP)* lines indicate that these cells have the morphological and migratory characteristics of myeloid progenitors (figure 1 supplement 2 – 4).

At 24 hpf, endocardial expression is observed in the linear heart tube of *Gt(endocard:egfp*) embryos (figure 1A). *Tg(fli1a:EGFP)* and *Tg(kdrl:Hsa.HRAS-mCherry)* are expressed in the endocardium but, in contrast, also show continuous expression with the adjacent vasculature (figure 1B). By 48 hpf, *Gt(endocard:egfp*) expression has expanded and is also observed in the developing vascular endothelium, indicating that this restriction from the vascular endothelium is transient. Additionally, at 24 and 48 hpf a number of GFP^+^/mCherry^-^ cells were observed surrounding the endocardial lining of the heart in *Gt(endocard:egfp*); *Tg(kdrl:Hsa.HRAS-mCherry)* but not *Tg(fli1a:EGFP)*; *Tg(kdrl:Hsa.HRAS-mCherry)* embryos (figure 1A). These GFP^+^ cells expressed noticeably weaker GFP levels than those observed in the endocardial layer and were confirmed to co-localise with the myocardial marker *Tg(myl7:mCherry-CaaX)* (figure 1 supplement 5), indicating *Gt(endocard:egfp*) expression in the myocardium at later stages.

To investigate the earliest expression of the *Gt(endocard:egfp*) line, *in situ* hybridisation was performed over a developmental time-course from the 5ss (11.5 hpf) through to 48 hpf examining *gfp* transcription (figure 2). *gfp* expression was first detected as early as the 8ss (13 hpf), this expression was observed in faint semilunar domains in lateral regions at the anterior of the embryo (figure 2) and were lost in a *npas4l/cloche* mutant (figure 2 supplement 1). These bilateral expression domains were observed more posteriorly at the 10ss. By the 15ss (16.5 hpf), this expression is found at the midline (figure 2) in the region of the endocardial core of the cardiac disc (figure 2), consistent with the observations made by examining fluorescence (figure 1). As *gfp* expression in this line precedes the reported expression of *nfatc1* in zebrafish (Wong et al. 2012), *nfatc1* expression was also analysed over the same time-course. Contrary to published reports, *nfatc1* expression was also found to initiate as early as the 8ss and was observed in strikingly similar domains to that of *gfp* expression in *Gt(endocard:egfp*) embryos (figure 2). Together, these results show that the endocardium begins to express multiple unique markers as early as the 8ss, much earlier than previously appreciated in zebrafish development. This expression is consistent with the observed fluorescence in the *Gt(endocard:egfp*) line and supports the hypothesis that the endocardial and vascular endothelial lineages diverge during early somitogenesis stages in the vertebrate embryo.

**Figure 2.**
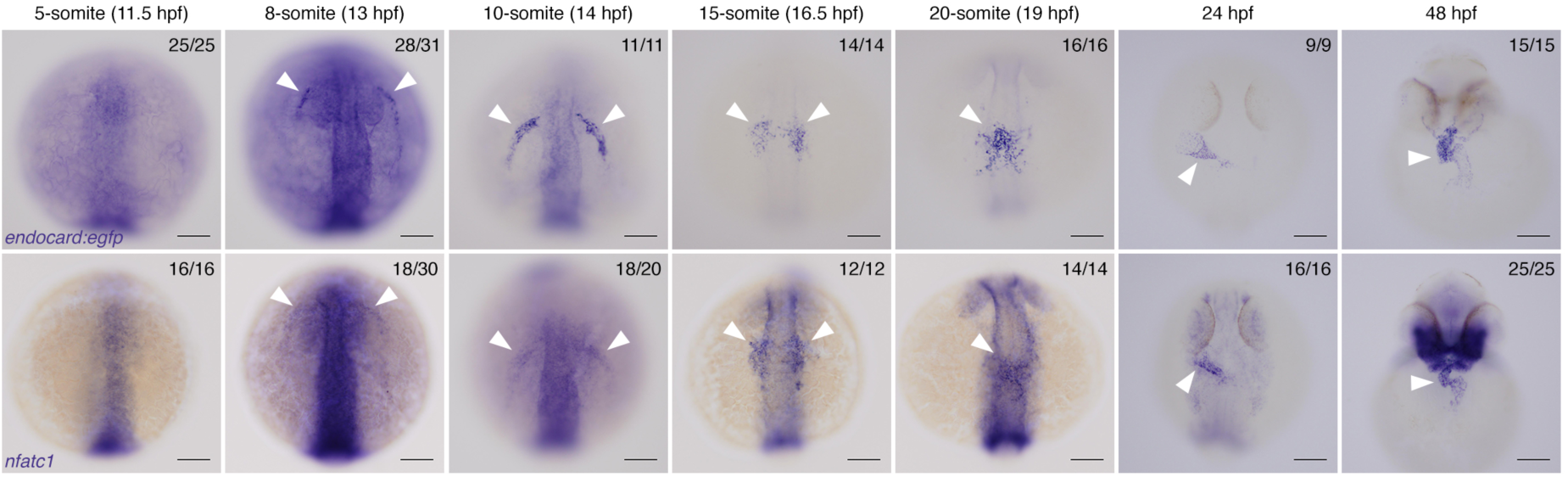
Restricted endocardial markers are first expressed at the 8ss in zebrafish embryos. *In situ* hybridisation for *gfp* expression in *Gt(endocard:egfp*) embryos and *nfatc1* expression from the 5ss (11.5 hpf) through to 48 hpf, showing expression emerging from 8ss onwards. White arrowheads show the expression domains corresponding to endocardial cells and their progenitors. Anterior to the top for 5ss – 24 hpf. Anterior views at 48 hpf, all other images show dorsal views. Scale bars represent 100 µm. The number of embryos matching the image is indicated in the top right of each image.

### Loss of myocardial and myeloid lineages do not alter early endocardial progenitor commitment

Having established the *Gt(endocard:egfp*) line as a marker of the endocardium, we sought to confirm its location in relation to other known tissues in the anterior lateral plate mesoderm using two-colour fluorescent *in situ* hybridisation.

Endocardial localisation was examined at the 15ss (16.5 hpf) using probes for *gfp* expression in *Gt(endocard:egfp*) embryos along with *spi1b* or *myl7* probes to label myeloid and myocardial populations, respectively (Yelon et al. 1999; Lieschke et al. 2002). Little overlap between *gfp* and *spi1b* expression was observed, with myeloid cells positioned anterior and medial to endocardial cells (figure 3 supplement 1). A lack of overlapping expression indicates that *gfp* is not actively transcribed in myeloid cells at the 15ss in *Gt(endocard:egfp*) embryos, contrasting the GFP fluorescence observed at the protein level (figure 1 supplement 2 - 3). This observation may be explained by perdurance of the GFP. Myocardial expression was located posterior to the endocardium at the 15ss, with no overlap observed between *gfp* and *myl7* expression (figure 3 supplement 1).

Previous studies have elegantly demonstrated that perturbation of fate-determining transcription factors can impact the expression domains of myocardial and myeloid populations across the ALPM (Lieschke et al. 2002; Keegan et al. 2005; Schoenebeck et al. 2007; Sumanas et al. 2008; Simões et al. 2011). However, without a unique marker of endocardial identity, this population is yet to be studied independent of the adjacent myeloid and vascular endothelial lineages.

To undertake this, we began by generating a range of loss-of-function models and examined the impact on endocardial development. Surprisingly, neither *nkx2.5*, *hand2* nor *tal1* loss-of-function were found to significantly affect the endocardial expression domain at the 14 or 16ss (16 and 17 hpf, respectively; figure 3 supplement 2 - 4). *hand2* mutants showed a drastic, but not complete, loss of the myocardial domain as previously reported (Yelon et al. 2000) without altering endocardial or myeloid lineages (figure 3 supplement 3). *tal1* knockdown caused a significant reduction of the myeloid domain (figure 3 supplement 4), consistent with previous reports (Shivdasani et al. 1995; Gering et al. 1998; Patterson et al. 2005). Interestingly, *tal1* knockdown increased the myocardial expression domain and expanded the length of this domain along the anterior-posterior axis without affecting the endocardial domain (figure 3 supplement 4 - 5).

Next, we examined the endocardial regulators *npas4l* and *etv2* in loss-of-function models. As expected, in *npas4l* mutants endocardial expression was completely lost at both 14 and 16ss when assessed by *in situ* hybridisation (figure 3A, E). The myeloid expression domain was also completely lost in mutants (figure 3C), consistent with previous reports (Lieschke et al. 2002), while the myocardial expression domains were expanded (figure 3G, figure 3 supplement 5), as described elsewhere (Schoenebeck et al. 2007).

**Figure 3.**
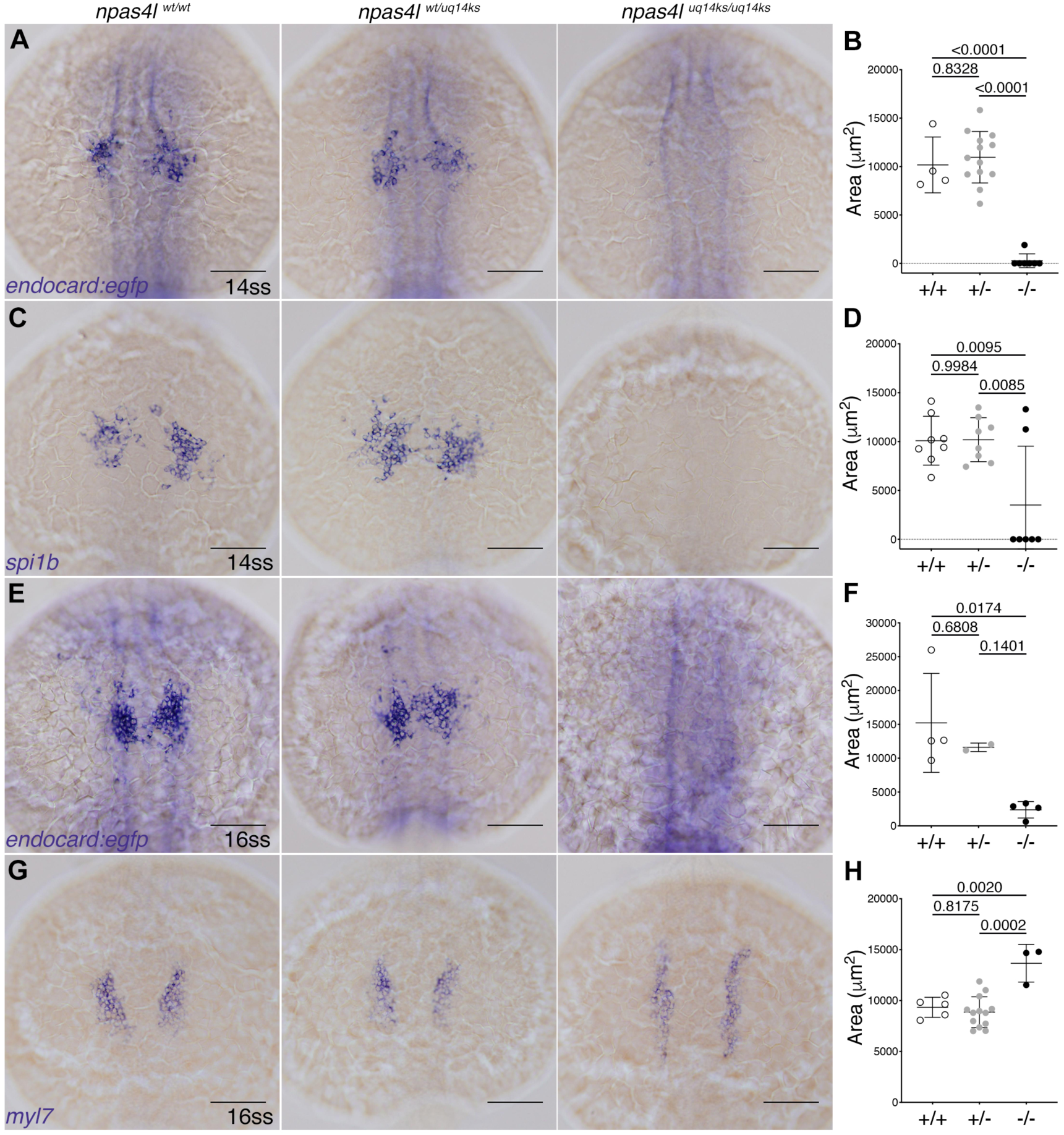
*Gt(endocard:egfp*) expression is lost in *cloche/npas4l* mutants. *In situ* hybridisation for **A.** *gfp* and **C.** *spi1b* expression at the 14ss and **E.** gfp and **G.** *myl7* expression at the 16ss in *npas4l^uq14ks^*mutants, heterozygotes and wildtype siblings. Quantification of expression domain is shown in **B**, **D**, **F** and **H**. Dorsal views, with anterior to the top. Scale bars represent 100 µm. P values are present in graphs.

In *etv2* mutants, endocardial expression was decreased as assessed by measuring the surface area of staining at the 14ss (figure 4A - B). Most notably however, the bilateral *gfp* expression in these mutants was extended along the anterior-posterior axis and localised to the lateral margins of the embryo rather than condensing and migrating to the midline as in wild-type siblings (figure 4A). By the 16ss, the *gfp* expression domain was markedly reduced (figure 4E - F). In contrast, *spi1b* expression was lost at the 14ss and *myl7* expression expanded at the 16ss (figure 4C, G).

**Figure 4.**
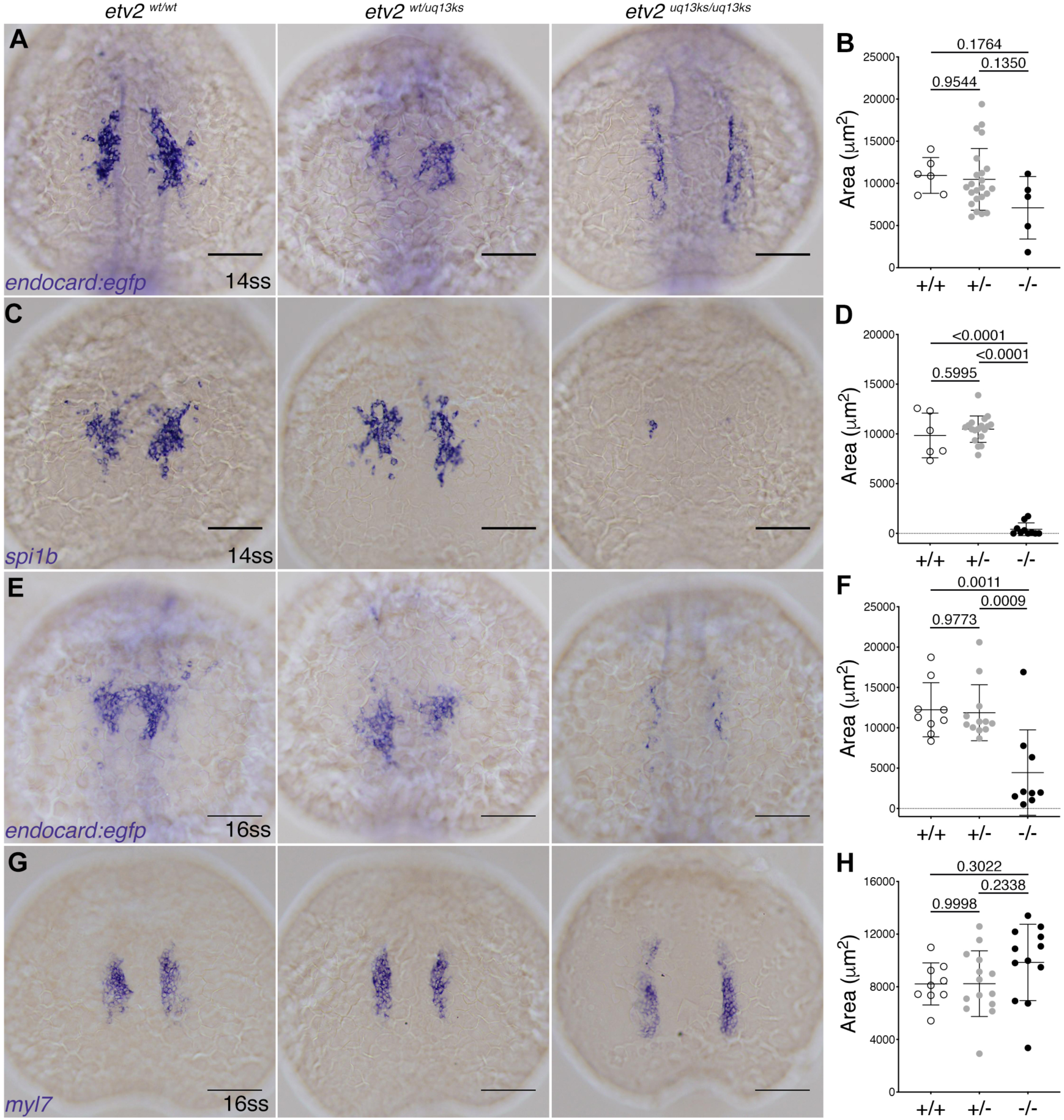
*Gt(endocard:egfp*) expression is reduced in *etv2* mutants. *In situ* hybridisation for **A.** *gfp* and **C.** *spi1b* expression at the 14ss and **E.** gfp and **G.** *myl7* expression at the 16ss in *etv2^uq13ks^* mutants, heterozygotes and wildtype siblings. Quantification of expression domain is shown in **B**, **D**, **F** and **H**. Dorsal views, with anterior to the top. Scale bars represent 100 µm. P values are present in graphs.

Together, these results show that the myocardial transcription factors *nkx2.5* and *hand2* are dispensable for early endocardial development. Similarly, *tal1* was also found to be dispensable, despite its role in maintaining endocardial identity during later development (Van Handel et al. 2012; Schumacher et al. 2013). Interestingly, while the myocardial domain was expanded in the absence of endocardial and myeloid domains, no reciprocal relationship was observed with a loss of myocardium in *hand2* mutants.

### BMP signalling is required for endocardial development

To identify regulators of endocardial development, a drug screen was performed using a number of inhibitors of major signalling pathways (figure 5 supplement 1). *Gt(endocard:egfp*) embryos were treated with chemical inhibitors, from the 5ss (11.5 hpf) until fixation at the 14 or 16ss. Fixed embryos were analysed by *in situ* hybridisation for markers of endocardial, myeloid and myocardial domains as described above.

Analysis at the 14ss showed slight decreases in the endocardial expression domain of embryos treated with VEGF, BMP and Wnt inhibitors relative to the DMSO control (with only the VEGF inhibitor treatment reaching statistical significance) (figure 5 supplement 1). By the 16ss, all three inhibitors were found to significantly reduce the endocardial expression domain relative to the DMSO control (figure 5 supplement 1). Inhibition of the Hedgehog pathway slightly increased the endocardial expression domain, however this was not statistically significant.

In zebrafish, VEGF signalling has previously been shown to act through Erk during early stages of angiogenesis (Shin, Beane, et al. 2016a; Shin, Male, et al. 2016b). To determine whether VEGF signalling may be acting through Erk in the endocardium, we investigated Erk activity using a phosphorylated Erk (pErk1/2) antibody. Surprisingly, immunofluorescence staining against pErk1/2 in *Gt(endocard:egfp*) or *Tg(fli1a:EGFP)* embryos at the 15ss did not show evidence of signalling in the endocardium at this stage (figure 5A - B). We simultaneously examined phosphorylated Smad1/5/8 (pSmad1/5/8), the downstream effector of BMP signalling (Derynck & Zhang 2003), and observed compelling pSmad1/5/8 activity in GFP^+^ cells in the endocardium at the 15ss using either *Gt(endocard:egfp*) or *Tg(fli1a:EGFP)* to co-label endocardium (figure 5C - D). Interestingly, in *Tg(fli1a:EGFP)* embryos pSmad1/5/8^+^/GFP^+^ cells did not co-localise with presumptive vascular endothelium (figure 5D), suggesting that BMP signalling is active in the endocardium but not the adjacent vascular endothelium of 15ss embryos.

**Figure 5.**
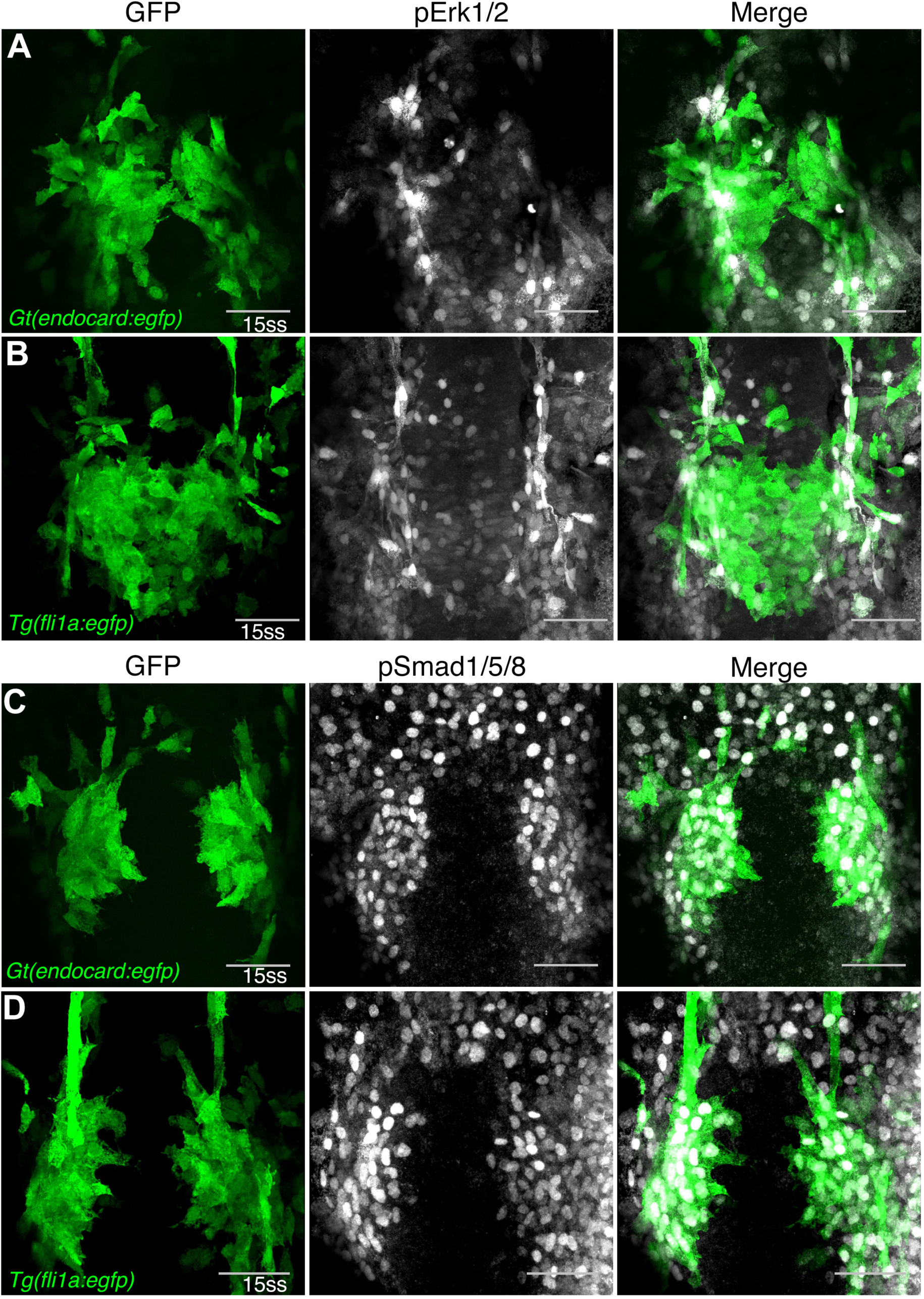
BMP is actively signalling in developing endocardial cells. Immunofluorescence staining for pErk1/2 in **A.** *Gt(endocard:egfp*) and **B.** *Tg(fli1a:egfp)* embryos at the 15ss shows minimal pErk1/2 signal in endocardial cells and high activity in adjacent vasculature. By contrast, pSmad1/5/8 in **C.** *Gt(endocard:egfp*) and **D.** *Tg(fli1a:egfp)* embryos at 15ss shows high pSmad1/5/8 activity in developing endocardial cell but minimal activity in adjacent vasculature. Dorsal views are shown with anterior to the top in all images. Scale bars represent 50 µm.

To examine the role of BMP signalling in endocardial development further, the *Gt(endocard:egfp*) line was crossed with the heat-shock-inducible transgenic lines, *Tg(hsp70l:bmp2b)* or *Tg(hsp70l:noggin3),* to either activate or inhibit the BMP pathway (Chocron et al. 2007). Embryos were heat-shocked at the tailbud stage (10 hpf) and fixed at the 14 or 16ss for analysis of myeloid, endocardial and myocardial lineages (figure 6 and figure 6 supplement 1). Inhibition of the BMP signalling pathway by global induction of *noggin3* expression resulted in a complete loss of endocardial (figure 6) and myeloid expression domains (figure 6 supplement 1) at the 14ss. The myocardial domain was also severely reduced in this context (figure 6 supplement 1). In the reciprocal experiment, activation of the BMP signalling pathway by global induction of *bmp2b* expression resulted in a significant expansion of the endocardial (figure 6) and myeloid (figure 6 supplement 1) expression domains but did not affect the myocardial domain (figure 6 supplement 1). Interestingly, activating BMP signalling not only increased the endocardial expression domain but also resulted in stronger staining compared with siblings lacking the heat-shock transgene (figure 6), indicating both an expansion of the endocardial domain and increased transcription within endocardial cells.

**Figure 6.**
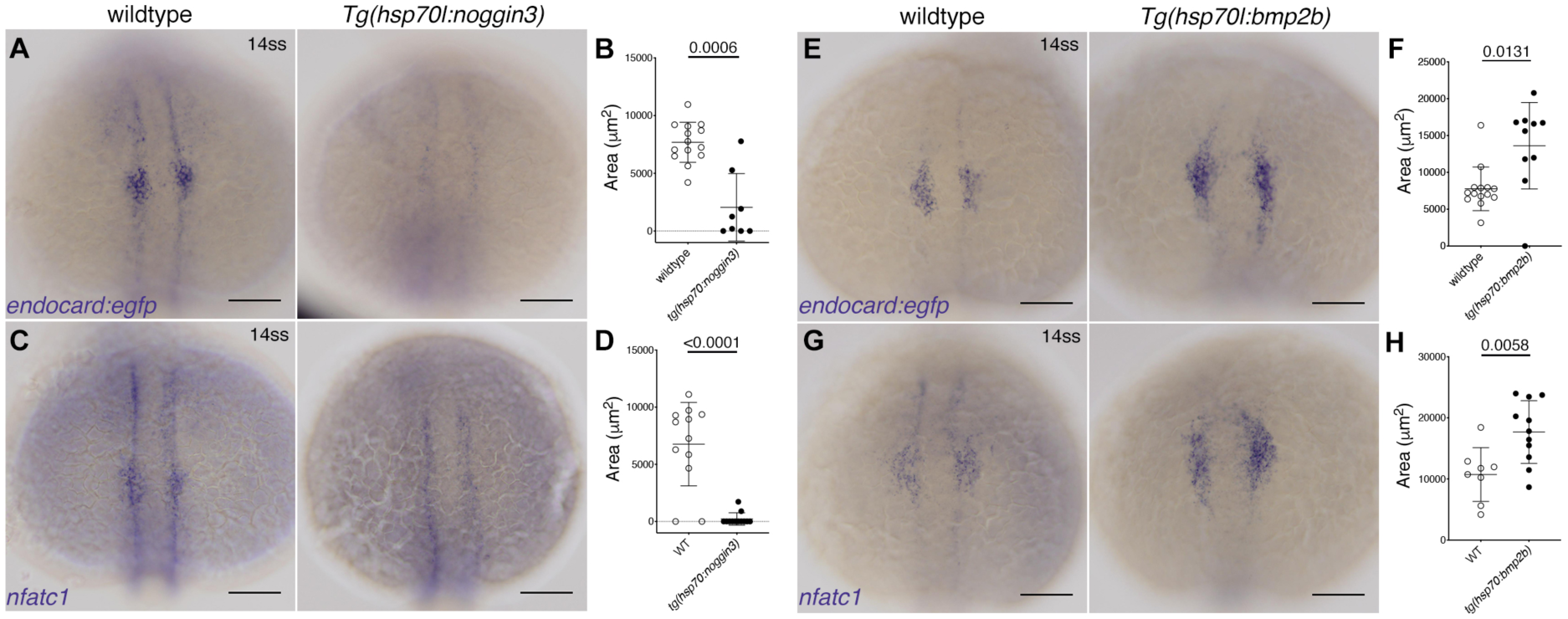
BMP signalling is required for endocardial development. *In situ* hybridisation for endocardial markers **A.** *gfp* (in *Gt(endocard:egfp)* embryos) and **C.** *nfatc1* in wildtype (transgenic negative siblings) or *Tg(hsp70l:noggin3)* embryos at the 14ss (heat-shocked performed at tailbud stage (10 hpf)) show significantly reduced staining of the endocardial domain upon inhibition of BMP signaling. **B** and **D.** Quantification of the area of expression shown in **A** and **C.** Reciprocally, wildtype sibling controls or *Tg(hsp70l:bmp2b)* embryos at 14ss (heat-shocked at tailbud) show increased staining area in **E.** *Gt(endocard:egfp)* embryos stained for *gfp* or **G.** embryos stained for *nfatc1*. **F** and **H**. Quantification of the area of expression shown in **E** and **G.** Dorsal views are shown with anterior to the top in all images. Scale bars represent 100 µm. P values are present in graphs.

Given the requirement of the endocardium for *etv2* activity (Figure 4) (Lee et al. 2008; Ferdous et al. 2009; Palencia-Desai et al. 2011), we sought to examine *etv2* expression under perturbation of BMP signalling. Activation of BMP signalling by heat-shock at the tailbud stage showed upregulated *etv2* expression in the anterior lateral plate mesoderm at the 14ss (figure 7). *etv2* expression in the posterior lateral plate mesoderm was also found to be upregulated in response to activation of the BMP signalling pathway (figure 7). Reciprocally, inhibition of BMP signalling by heat-shock at the tailbud stage altered the patterning of the expression domain of *etv2* in both the anterior and posterior lateral plate mesoderm (figure 7).

**Figure 7.**
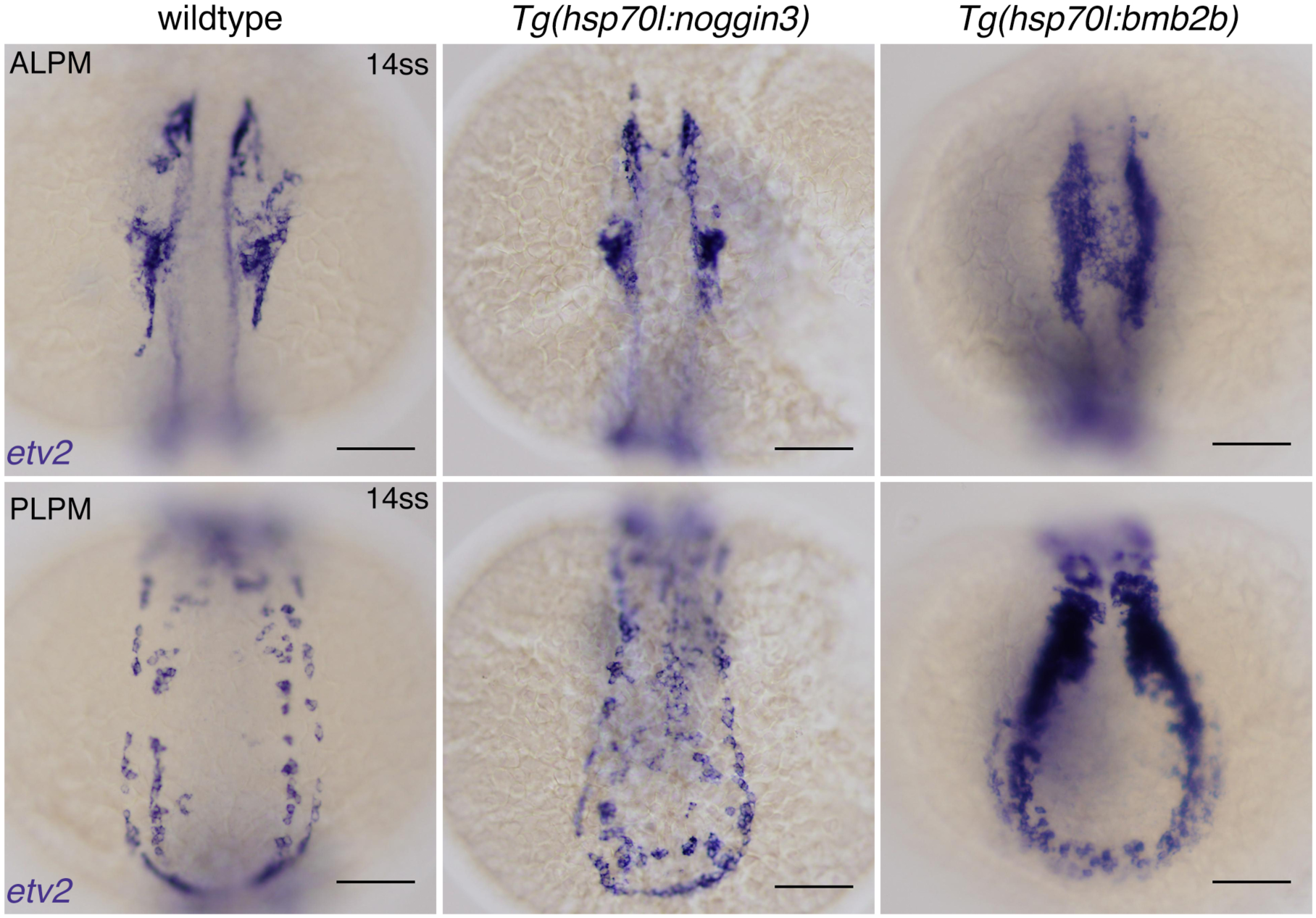
*etv2* expression is altered by BMP signaling modulation. *In situ* hybridisation for *etv2* expression in wildtype sibling controls, *Tg(hsp70l:noggin3)* or *Tg(hsp70l:bmp2b)* embryos. The *etv2* expression domain is altered with inhibition of BMP signaling and expanded and upregulated with overactivation of BMP signaling. Dorsal views are shown with anterior to the top in all images. 9/9 *Tg(hsp70l:noggin3)* embryos matched the expression pattern shown while 11/14 WT embryos matched the WT pattern shown. 13/13 *Tg(hsp70l:bmp2b)* embryos matched the expression pattern shown while 10/10 WT embryos matched the WT pattern shown Scale bars represent 100 µm. ALPM - anterior lateral plate mesoderm, PLPM - posterior lateral plate mesoderm.

Together, these results show that BMP signalling plays a crucial role in early endocardial development and is essential for correct *etv2* expression, a gene essential for endocardial development. Both BMP inhibition and *etv2* loss-of-function result in a loss of endocardium, suggesting that BMP signalling acts upstream of *etv2* to induce endocardial development.

## Discussion

The endocardium is a unique invention of the vertebrate taxa (Pérez-Pomares et al. 2009). Although the endocardium has many molecular similarities to the adjacent vascular endothelium, there are distinct physiological functions of these tissues (Harris & Black 2010). What distinguishes these lineages on a molecular level remains an important, unanswered question in cardiovascular biology. Using zebrafish as a model for cardiac development, we have identified the *Gt(endocard:egfp*) gene-trap transgenic line as an endocardial marker and used this to profile the molecular signals required for endocardial differentiation.

Characterisation of the *Gt(endocard:egfp*) line showed that this marker is uniquely expressed in the progenitors of the endocardium as early as the 8ss (13 hpf) - 9 hours prior to the reported initiation of expression of the endocardial specific marker, *nfatc1* (Wong et al. 2012). By re-examining *nfatc1* expression in zebrafish embryos it was found that, contrary to previous reports, *nfatc1* expression initiates at the same stage as the *Gt(endocard:egfp*) marker. Although the faint expression of *nfatc1* prevents co-localisation by fluorescent *in situ* hybridisation, these markers are expressed in a strikingly similar spatiotemporal pattern. This pattern is consistent with *npas4l* expression, the earliest known regulator of endocardial fate, during early somitogenesis stages (Reischauer et al. 2016) and the observed expression domain is lost in the *npas4l^uq12ks^* mutant. These results demonstrate that the 8ss marks the emergence of definitive markers of the endocardium in the anterior lateral plate mesoderm.

This 8ss is intriguing as *tal1* knockdown embryos also develop gene expression defects at this time (Patterson et al. 2005). In *tal1* morphants, the mesodermal markers *hhex* and *draculin* are substantially reduced from the 7ss, despite initiating expression normally at earlier stages (Patterson et al. 2005). Although our results did not find any association between *tal1* loss-of-function and early endocardial development, *tal1* has been shown to be required in endocardial development at later stages in zebrafish and mice (Bussmann et al. 2007; Van Handel et al. 2012; Schumacher et al. 2013).

Furthermore, both *hhex* and *draculin* are associated with cardiovascular development (Herbomel et al. 1999; Liao et al. 2000; Mosimann et al. 2015; Gauvrit et al. 2018). These results point to the 7/8ss as a critical phase in the establishment of cardiovascular lineages in zebrafish development.

Using *gfp* expression in the *Gt(endocard:egfp*) line as a proxy for endocardial gene expression, the relationship between the endocardial, myeloid and myocardial lineages was probed in a number of loss-of-function models. In these models, endocardial and myeloid lineages were regulated in kind in most models examined, suggesting a close association between the development of the endocardial and myeloid lineages in the zebrafish. When endocardial and/or myeloid lineages were lost, there was a resulting increase in the myocardial expression domain, as has previously been reported for *cloche* mutants (Schoenebeck et al. 2007) and *etv2* morphants (Palencia-Desai et al. 2011). However, no evidence for a reciprocal increase in the endocardial or myeloid lineages was seen when the myocardial domain was ablated in *hand2* mutants.

While FGF signalling has previously been shown to have reciprocal effects on haemangioblast and myocardial lineages, these effects were most pronounced at the blastula stage (Simões et al. 2011). The results from our experiments, combined with the previous work described above, suggest that there is an early step directing commitment of cardiovascular progenitor cells to the haemangioblast or myocardial lineages. After becoming committed to the haemangioblast programme, additional signals are required to maintain the differentiation of these lineages and repress myocardial development. In the absence of these additional signals, mesodermal cells fail to continue developing along the haemangioblast lineage and default to the myocardial program. In support of this theory, scRNA-seq analysis of mesodermal cells in *Tal1* mutant mouse embryos has shown that in the absence of *Tal1*, haemogenic and endocardial progenitors revert to a myocardial lineage (Van Handel et al. 2012; Scialdone et al. 2016). Moreover, this was not found to be through direct regulation of the myocardial program by *Tal1* (Scialdone et al. 2016), suggesting that establishment of endocardial identity restrains the myocardial program and simply halting endocardial differentiation is sufficient to initiate myocardial development.

Additional analysis of the endocardial, myeloid and myocardial lineages using inhibitors and genetic tools, as well as the analysis of downstream effectors of signalling pathways, identified the BMP signalling pathway as playing a vital role in the development of the endocardial lineage. Interestingly, inhibition of the BMP signalling pathway in zebrafish by drug treatment from the 5ss had mild effects on the endocardial and myeloid lineage compared to *noggin3* over-expression from the tailbud stage. Similarly, while no significant affect was observed on the myocardium by drug treatment from the 5ss, *noggin3* over-expression at the tailbud stage resulted in a drastic loss of myocardium. This suggests the early somitogenesis stages mark a crucial window when cardiovascular progenitors transition from an uncommitted to committed state. Furthermore, our results show that BMP signalling is both necessary and sufficient for the development of the endocardial and myeloid lineages, and necessary for the development of the myocardial lineage.

The requirement for BMP signalling in endocardial development appears to be conserved in mice, as deletion of the BMP receptor, Alk2, in endothelium but not myocardium results in endocardial defects (Wang et al. 2005). Similarly, activation of *Noggin* in the endocardium using a constitutively actively *Nfatc1^Cre^*line results in endocardial-specific defects (Snider et al. 2014). Interestingly, BMP signalling is also sufficient for the maintenance of endocardial identity: zebrafish embryos that are myocardial-deficient fail to express *nfatc1*, demonstrating myocardium is necessary for the maintenance of endocardial identity. In this context, activation of BMP signalling at 16ss was capable of rescuing *nfatc1* expression at 24 and 46 hpf (Palencia-Desai et al. 2015), suggesting that the endocardium is dependent upon BMP signalling, potentially derived from the myocardium at these later stages and from an unknown source at earlier stages. These results support a conserved and ongoing role for BMP signalling in endocardial development.

Here, we report that the genetic activation or inhibition of the BMP signalling pathway modulates *etv2* expression. As *etv2* loss-of-function was also found to result in a loss of endocardial progenitors, these results have led to a proposed model whereby BMP signalling controls *etv2* expression, which in turn directs commitment to the endocardial and myeloid lineages. A number of regulators of *etv2* expression have been identified in zebrafish (Veldman & Lin 2012; Schupp et al. 2014), as well as in mice (Rasmussen et al. 2012; Koyano-Nakagawa & Garry 2017). In support of our hypothesis, BMP signalling has also been shown to regulate *etv2* expression through controlling neuromuscular progenitor fate in the zebrafish trunk (Row et al. 2018). Furthermore, two conserved regions in the upstream promoter of murine *Etv2* have been identified as containing Smad-binding motifs (Shi et al. 2015). These results suggest that BMP signalling may control *etv2* expression by direct interaction of Smads with the *etv2* promoter. However, it was recently reported that *etv2* is directly regulated by *npas4l* binding to the *etv2* locus (Marass et al. 2019). As the timing of *npas4l* expression and BMP activity are coincident in the embryo, it remains to be determined whether *npas4l* and Smads function co-operatively, or in parallel, to control the expression of *etv2*.

Together, these results represent the identification of a novel tool for investigating early endocardial development and report defined endocardial expression domains prior to linear heart tube formation in zebrafish. Furthermore, through analysing the endocardial expression domain we have identified BMP signalling as a critical regulator of early endocardial development. Based on these results, we propose a model for endocardial development whereby BMP signalling acts upstream of the transcription factor *etv2* to regulate the emergence of endocardial progenitors from the anterior lateral plate mesoderm in zebrafish.

## Materials and Methods

### Zebrafish lines

All zebrafish strains were maintained and animal work performed in accordance with the guidelines of the animal ethics committee at The University of Queensland, Australia. The previously published transgenic lines used in this study are *Gt(SAGFF27C); Tg(4xUAS:GFP),* referred to as *Gt(endocard:egfp*) in this text (Bussmann et al. 2010), *Tg(kdrl:Hsa.HRAS-mCherry)^s916^*(Hogan et al. 2009), *Tg(fli1a:EGFP)^y1^* (Lawson & Weinstein 2002), *Tg(hsp70l:noggin3)^fr14^* and *Tg(hsp70l:bmp2b)^fr13^*(Chocron et al. 2007).

### Morpholino oligonucleotide (MO) reagents

All MOs were ordered from Genetools, LLC. The MO sequences and concentrations used are as follows: 0.8 pmol *tal1*/*scl* - AATGCTCTTACCATCGTTGATTTCA (Dooley et al. 2005), 1 pmol *gata1a* - CTGCAAGTGTAGTATTGAAGATGTC (Galloway et al. 2005), 1.77 pmol *spi1b* - GATATACTGATACTCCATTGGTGGT (Rhodes et al. 2005).

### CRISPR/Cas9 mutagenesis

To generate the mutant lines used in this study CRISPR/Cas9 mutagenesis was performed as described previously (Gagnon et al. 2014; Capon et al. 2016). Embryos and fish were screened for indels by HRMA (Dahlem et al. 2012) and carriers sequenced to identify frame-shift mutations that truncate the protein. Specific details of each of the mutants generated for this study are below.

The *nkx2.5* mutant allele carries a 28bp deletion in exon one of the *nkx2.5* gene and is referred to as the *nkx2.5^uq15ks^*allele. This mutated allele results in a predicted protein product lacking the DNA binding homeobox domain. An incross of *nkx2.5^uq15ks^* heterozygous fish produces a mutant phenotype in 25% of progeny at 72 hpf consisting of a cardiac oedema and dilated atrium similar as previous reported (Targoff et al. 2008; Targoff et al. 2013)

The *hand2* mutant allele carries a 4bp deletion in the first exon of the *hand2* gene and is referred to as the *hand2^uq16ks^* allele. The truncated protein product of this mutant allele lacks the DNA binding bHLH domain. Incrossing fish carrying the *hand2^uq16ks^* allele results in a mutant phenotype in 25% of the progeny in which the affected embryos completely lack a beating heart and have markedly smaller fin buds as previously described (Yelon et al. 2000).

The *npas4l* mutant allele carries a 7bp deletion in the second exon of the *npas4l* gene and is referred to as the *npas4l^uq14ks^* allele. The predicted protein product of this truncated allele partially retains the DNA-binding bHLH domain, but lacks the two PAS domains as well as the three transcriptional activation domains. An incross of fish carrying the *npas4l^uq14ks^*allele produces a mutant phenotype in 25% of the progeny, consisting of a bell-shaped heart and lacking the endocardial layer at 48 hpf as previously described for the *cloche* mutant (Stainier et al. 1995).

The *etv2* mutant allele has a 23bp deletion in the exon five of the *etv2* gene and is referred to as the *etv2^uq13ks^*allele. This truncated allele completely lacks the DNA-binding ETS domain. An incross of fish heterozygous for the *etv2^uq13ks^*allele produces a mutant phenotype in 25% of progeny. This phenotype is characterised by a lack of blood circulation, as previously reported (Pham et al. 2007) as well as a collapsed heart at 48 hpf. Interestingly, the cardiac defect at 48 hpf was variable with the most severely affected mutant embryos lacking the majority of the endocardium.

### *In situ* hybridisation

In situ hybridisation was performed as previously described (C. Thisse & B. Thisse 2008). All RNA probes were transcribed from plasmids. Plasmids containing the *spi1b* and *gfp* coding sequences were a kind gift from the laboratory of Ben Hogan. To generate the *etv2* plasmid used in this study, pCS2+ vector was prepared by digesting empty vector with BamHI and XbaI. The *etv2* cDNA sequence was amplified by PCR using the primer sequences below, the PCR product was purified and inserted into the digested pCS2+ vector using circular polymerase extension cloning (Quan & J. Tian 2011).

*etv2*_cDNA_F:

AAGCTACTTGTTCTTTTTGCAGGATCTGTCAAAACCCCTGATATAGTG

*etv2*_cDNA_R:

TGGATCTACGTAATACGACTCACTATAGTTCTAGCAATCTGCTGCAAAGTCC

### Fluorescent *in situ* hybridisation

Fluorescent *in situ* hybridisation was performed as previously described (Clay & Ramakrishnan 2005; Brend & Holley 2009; Baek et al. 2019) with minor alterations. In brief, embryos were dechorionated and fixed at the desired stage. RNA probes were synthesised with DIG or FLU RNA labelling mix (Roche). Fixed embryos were permeabilised with proteinase K (Invitrogen) and hybridised with 1ng/µL RNA probe in hybridisation buffer overnight at 70°C. After hybridisation, embryos were washed, blocked with western blocking reagent (Roche) and incubated with anti-dig or anti-flu, POD antibodies (Roche) in western blocking reagent overnight at 4°C. After further washes, staining was performed using the tyramide signalling amplification kit (Perkin Elmer) for 2 hours at 37°C. Following staining embryos were fixed at 4°C overnight and mounted for imaging.

### Heat-shock treatment

Heat-shock was performed at 39°C for 30 minutes in pre-warmed medium. Following heat-shock, embryos were returned to a 28.5°C incubator until fixation.

### Antibody staining

Phospho-protein staining was performed as previously described (Okuda et al. 2018) using the following antibodies: GFP (abcam cat. no. ab13970), pErk1/2 (cell signaling technologies cat. no. 4370), pSmad1/5/8 (cell signaling technologies cat. no. 13820).

### Drug treatments

Chemical inhibitors of the following signalling pathways were ordered from Merck: cyclopamine - hedgehog inhibitor (cat. no. C4116), DAPT - Notch inhibitor (cat. no. D5942), DMH1 - BMP inhibitor (cat. no. D8946), IWR1 - Wnt inhibitor (cat. no. I0161), SU5416 - VEGF inhibitor (cat. no. S8442). These inhibitors were dissolved in DMSO, aliquoted, and stored at −20°C. For treatments, inhibitors were dissolved in E3 medium to the indicated concentration below, added to embryos at a ratio of 15 embryos/mL and incubated in the dark at 28.5°C until fixation.

| Drug name | Signalling pathway inhibited | Working concentration |
| --- | --- | --- |
| Cyclopamine | Hedgehog | 50 µM |
| DAPT | Notch | 100 µM |
| DMH1 | BMP | 1 µM |
| IWR1 | Wnt | 10 µM |
| SU5416 | VEGF | 1 µM |

### Imaging

Embryos from *in situ* hybridisation were dehydrated, cleared in murray’s solution (a 2:1 ratio of benzyl benzoate:benzyl alcohol) and imaged using an Olympus BX51 Microscope with Olympus DP70 CCD camera.

For wide-field bright-field and fluorescence imaging, embyros were mouted in 3% methyl cellulose (Sigma, cat. no. M0387) and imaged using a Leica M165 FC stereo microscope with a DFC425 C camera.

All confocal imaging was performed on a Zeiss LSM 710 FCS confocal microscope. Live embryos were mounted for confocal imaging using 0.5-1% low-melting agarose (Sigma, cat. no. A9414) in glass-bottom petri dishes. Fixed embryos for confocal imaging were de-yolked using lash tools and mounted in vectashield with DAPI (Vector laboratories, cat. no. H-1200) on glass-slides with coverslips.

All images were processed using FIJI (Schindelin et al. 2012) and/or Imaris software.

### Statistical testing

A student’s t-test with Welch’s correction was performed using Prism software. For data-sets containing 3 or more conditions, one-way ANOVA testing with Tukey’s post hoc test for multiple comparisons was performed using Prism software.

## Acknowledgements

We acknowledge members of the Smith laboratory for useful discussions and Ben Hogan and lab members for discussions and sharing of reagents. We thank the UQBR aquatics team for animal husbandry. Confocal microscopy was performed at the Australian Cancer Research Foundation’s Cancer Ultrastructure and Function Facility at the IMB.

## Funding

SJC was supported by an Australian Government Australian Postgraduate Award (APA) and KAS was part-funded by an Australian Research Council (ARC) Future Fellowship (FT110100496). The research was funded by an ARC Discovery Project grant (DP170101217).

## Competing interests

The authors declare that no competing interests exist.

## Supplemenatary material

**Figure 1 supplement 1.**
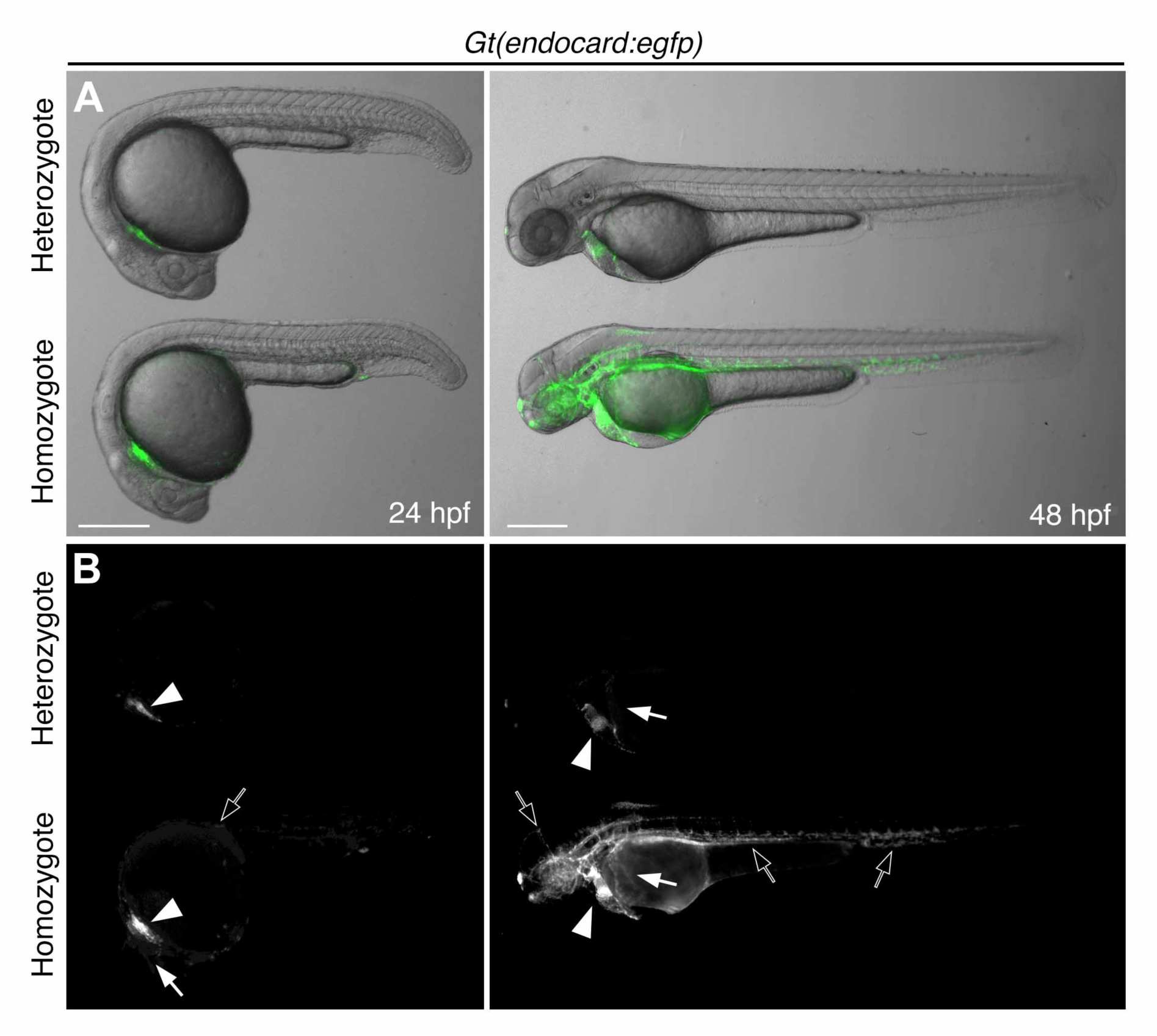
Live lateral view imaging of *Gt(endocard:egfp)* embryos at 24 and 48 hpf, **A.** with and **B.** without brightfield. Each panel shows a homozygous and heterozygous embryo imaged side-by-side in a single image. A low-GFP fluorescing embryo, heterozygous for both *Gt(SAGFF27C)* and *Tg(UAS:GFP)* transgenes, is shown above a high-GFP fluorescing embryo, homozygous for both *Gt(SAGFF27C)* and *Tg(UAS:GFP)* transgenes. Composite images are shown in **A** while GFP only is shown in **B**. Scale bars represent 500 µm.

**Figure 1 supplement 2.** Live, time-lapse imaging of a representative *Gt(endocard:egfp)* embryo beginning at the 11-somite stage. Confocal z-stacks were acquired at 5-minute intervals over a 4-hour period. Dorsal views are shown with anterior to the top. The bilateral populations of endocardial progenitors are observed emerging in the lateral margins of the ALPM before rapidly migrating to the midline to form the endocardial core of the cardiac disc. A small number of myeloid cells are observed emerging from the ALPM simultaneously with the endocardial progenitors and migrating away from the midline.

**Figure 1 supplement 3.** Live, time-lapse imaging of a representative *Tg(fli1a:egfp)* embryo beginning at the 12-somite stage. Confocal z-stacks were acquired at 5 minute intervals over a 4 hour period. Dorsal views are shown with anterior to the top. Bilateral populations of endocardial progenitors are observed migrating to the midline to form the endocardial core of the cardiac disc. Compared with *Gt(endocard:egfp)* embryos, a large number of myeloid cells are observed emerging from the ALPM and migrating away from the midline. A number of GFP positive cells are also observed in the lateral margins forming the cranial vascular endothelium while endothelial progenitors from the trunk are observed coming into frame as the migrate anteriorly.

**Figure 1 supplement 4.**
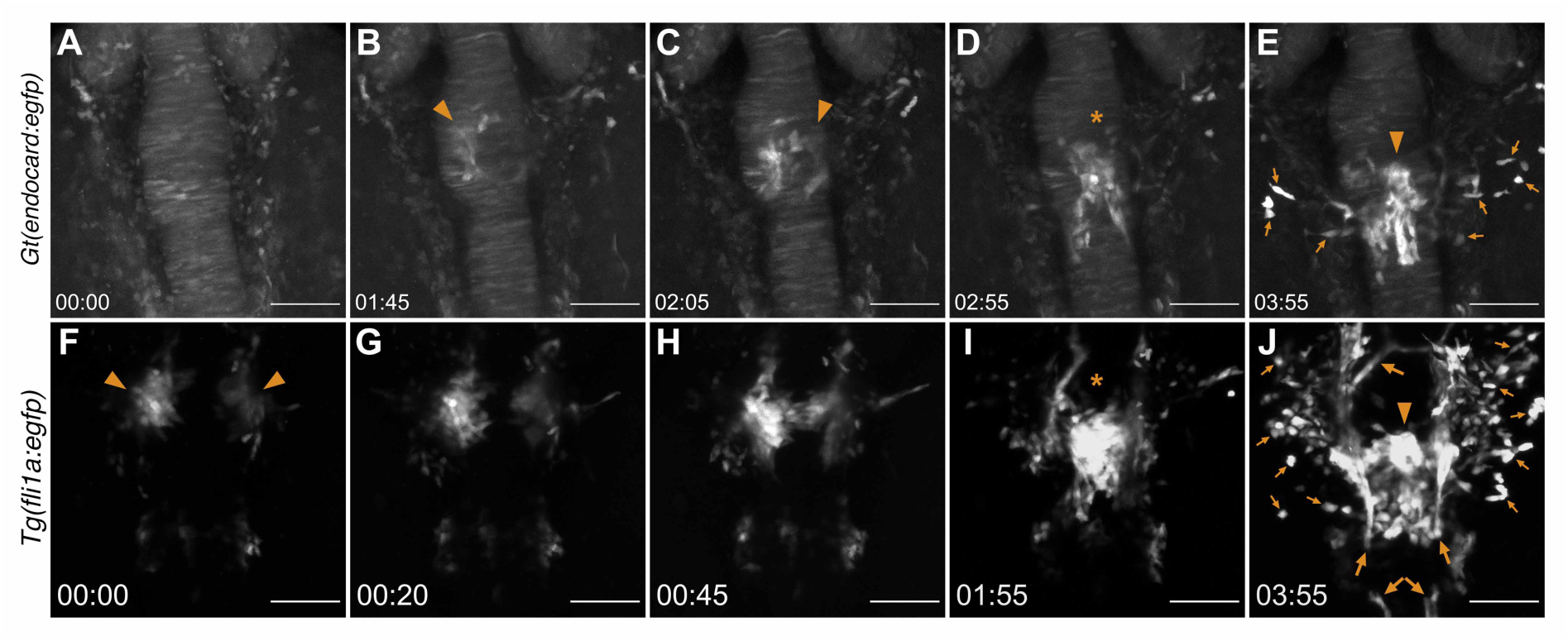
Still images from a representative time-lapse imaging on a *Tg(fli1a:EGFP)* embryo from the 12-somite stage (15 hpf) (A - E) and a representative *Gt(endocard:egfp)* embryo from the 11-somite stage (14.5 hpf) (F - J). Z-stacks were acquired at 5 minute intervals over a 4 hour period. Dorsal view is shown with the anterior to the top. Orange arrowheads point to emerging endocardial progenitors. Orange arrows point to migratory myeloid and vascular endothelial cells. Orange asterisk shows the endocardial progenitors after merging in the midline.

**Figure 1 supplement 5.**
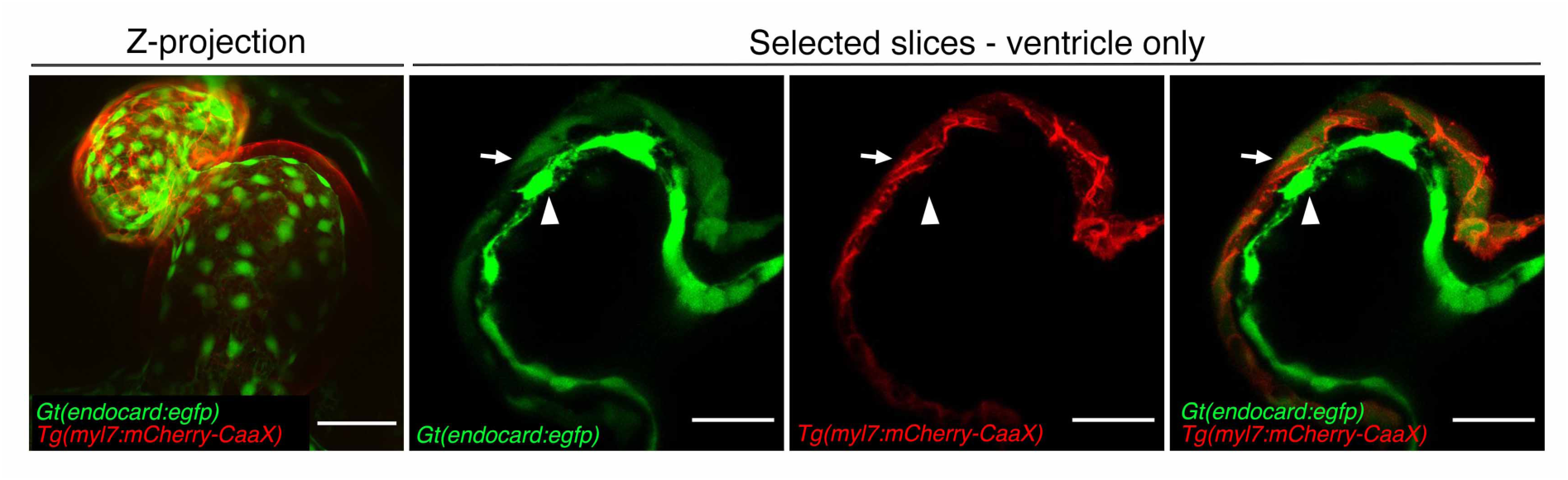
Live imaging of *Gt(endocard:egfp)*; *Tg(myl7:mCherry-CaaX)* embryos at 48 hpf. A Z-projection of the full heart is shown. Selected slices of the ventricle are also shown. All images are ventral views with anterior to the top. Scale bars represent 50 µm (in Z-projection) and 25 µm (in slices). White arrowheads label endocardium, white arrows label myocardium.

**Figure 2 supplement 1.**
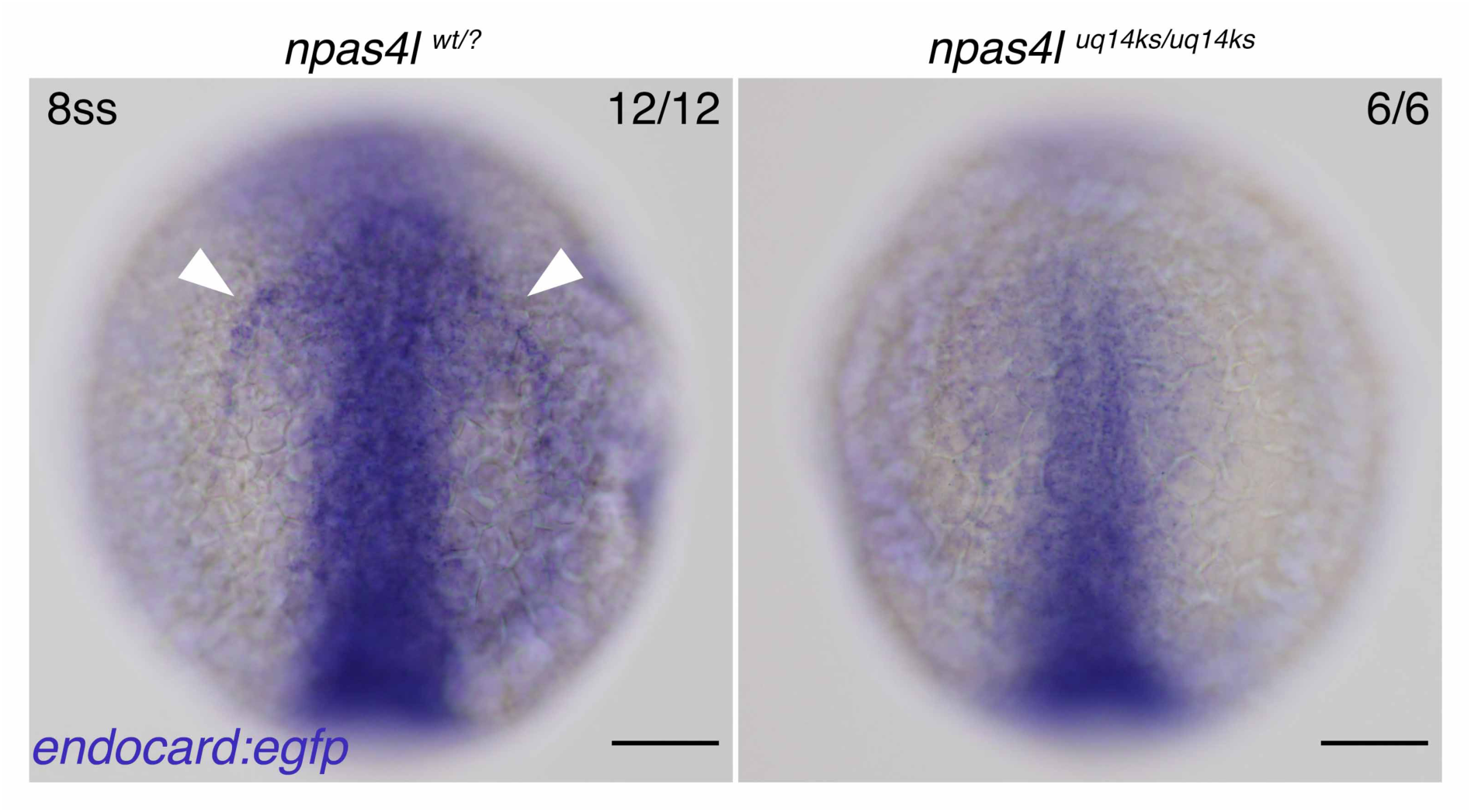
*In situ* hybridisation for endocardial *gfp* expression in *Gt(endocard:egfp)*; *npas4l* WT sibling and homozygous mutant embryos at the 8ss (13 hpf). Dorsal views in all images. Scale bars represent 100 µm. The number of embryos matching the image is indicated in the top right of each image.

**Figure 3 supplement 1.**
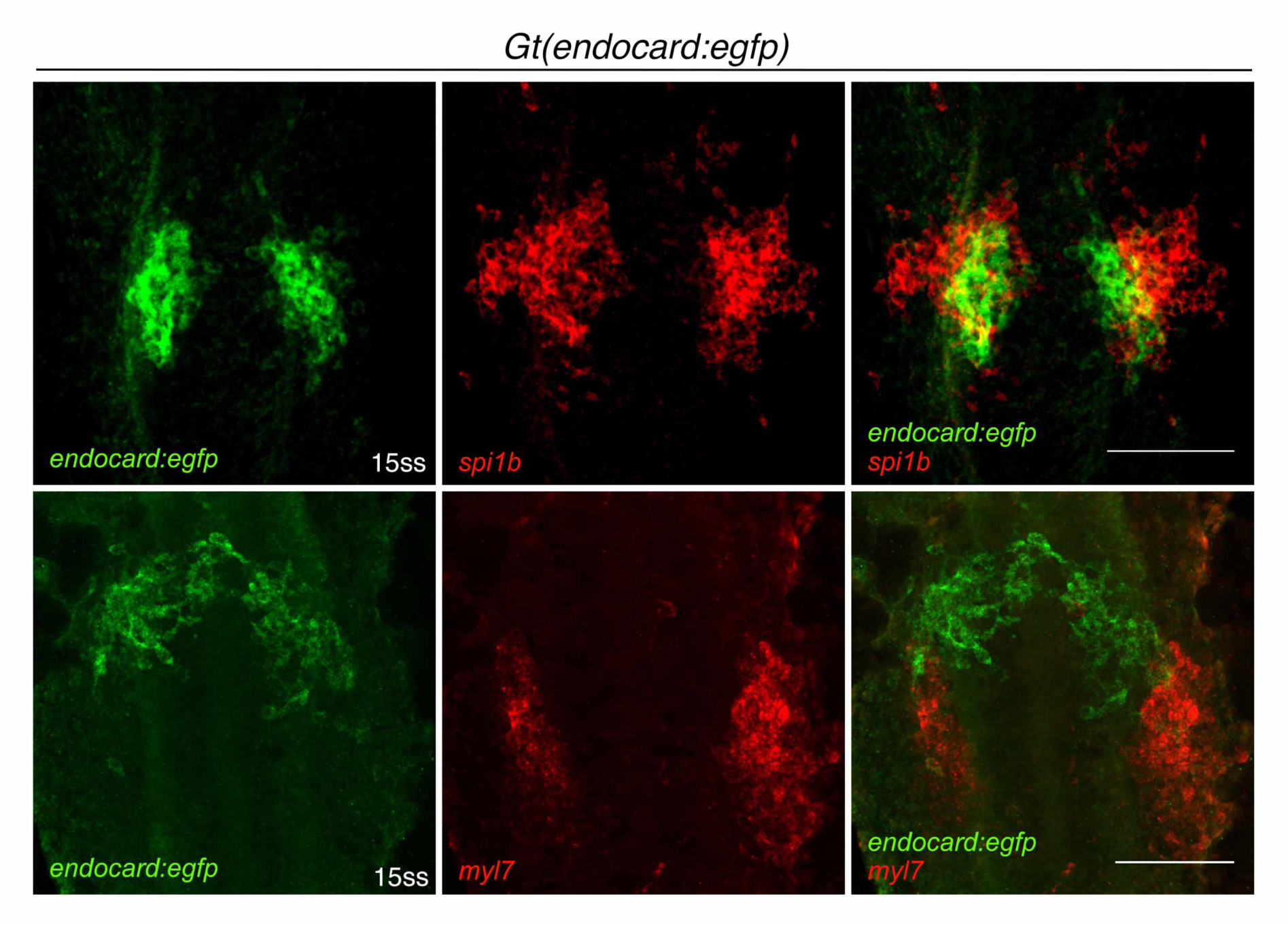
Fluorescent *in situ* hybridisation for *gfp* expression in *Gt(endocard:egfp)* embryos in combination with expression of the myeloid marker, *spi1b* (n=3), or the myocardial marker, *myl7* (n=3). Embryos were collected at the 15ss (16.5 hpf). Dorsal views are shown with anterior to the top in all images. Scale bars represent 100 µm.

**Figure 3 supplement 2.**
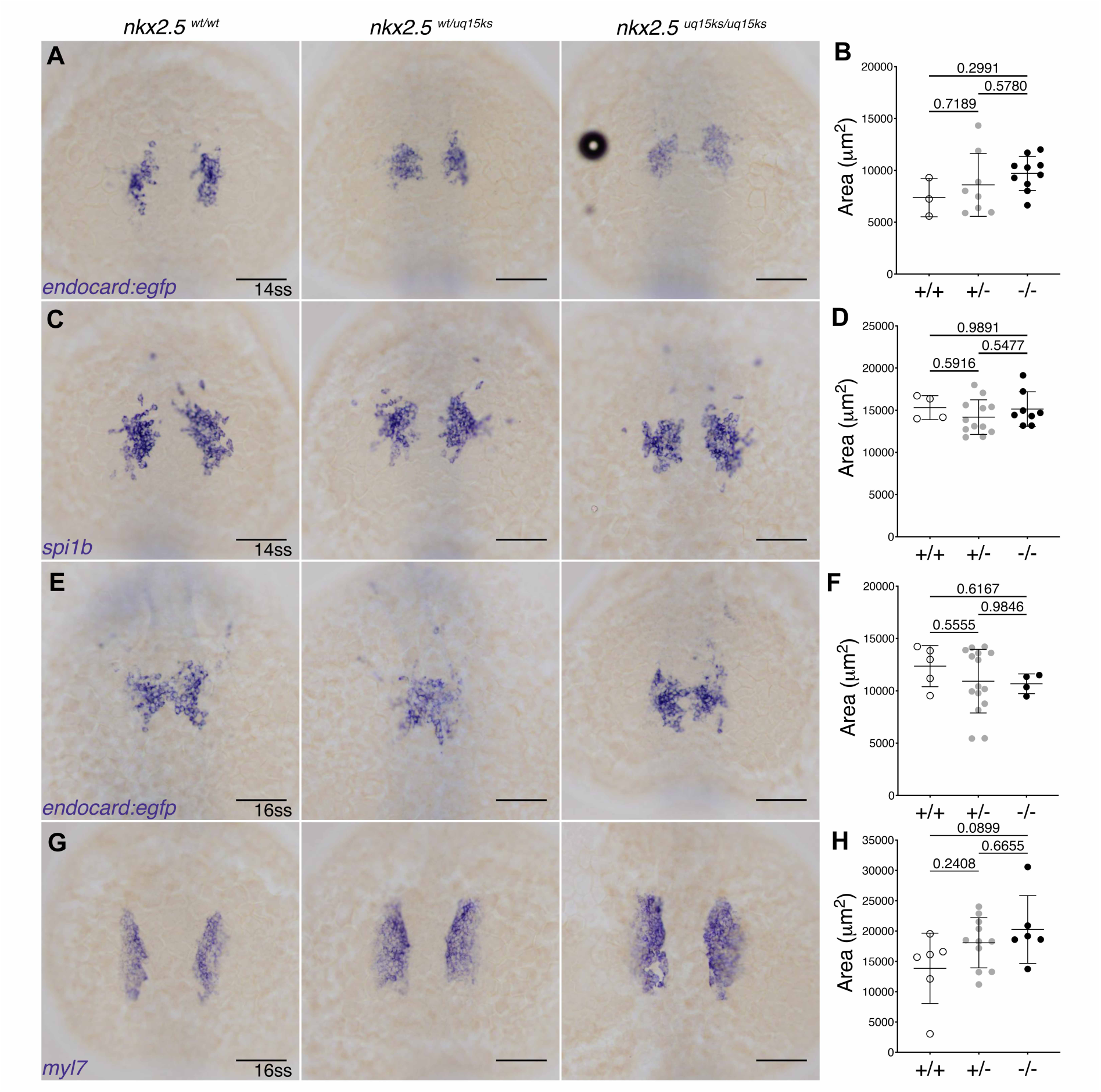
*In situ* hybridisation for endocardial *gfp* (A,E), myeloid, *spi1b* (C), and myocardial, *myl7* (G), markers in *nkx2.5^uq15ks^* mutants and siblings. Quantification of the expression is shown in B, D, F and H. Dorsal views are shown with anterior to the top in all images. Scale bars represent 100 µm. P values are indicated in graphs.

**Figure 3 supplement 3.**
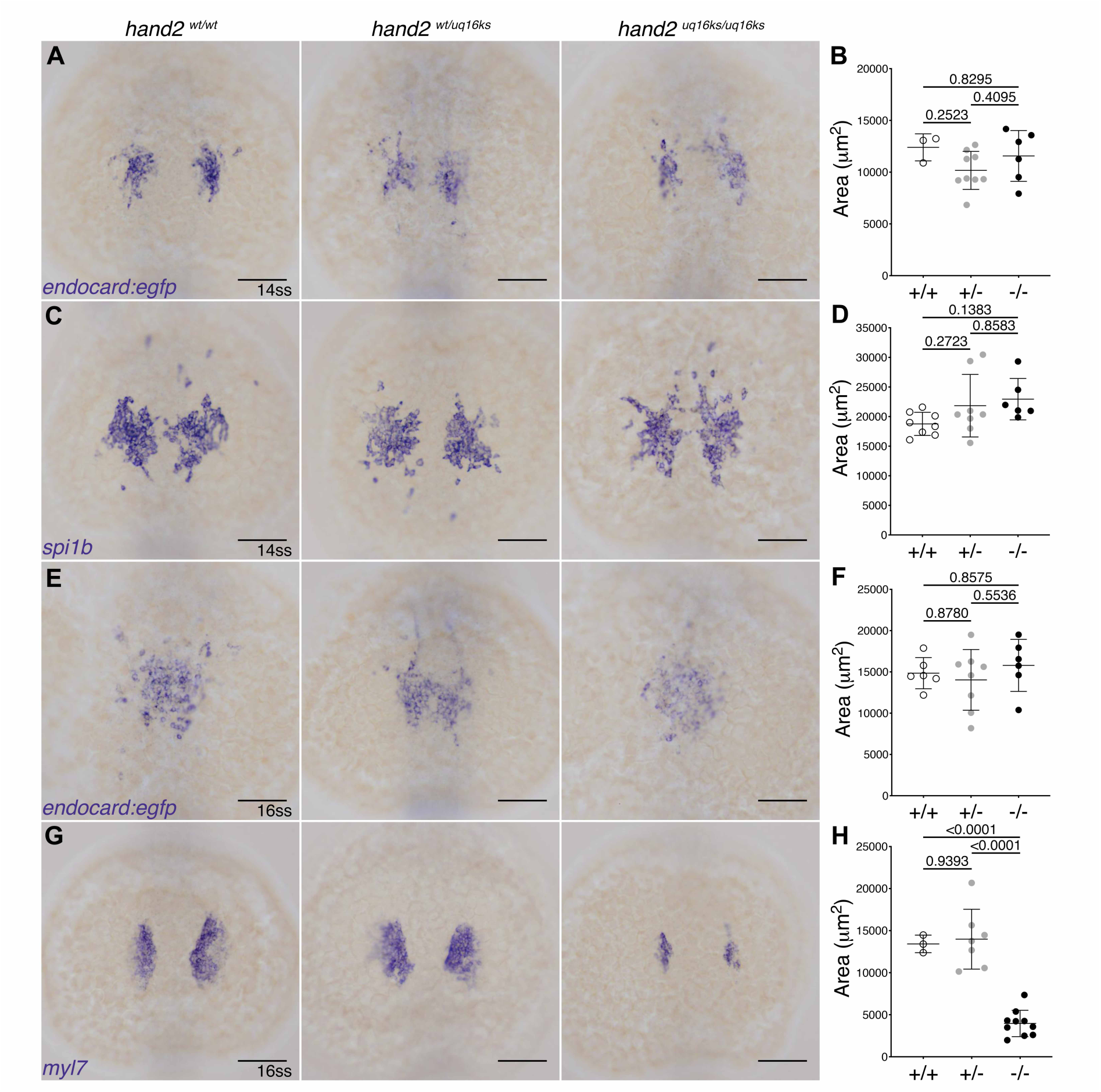
*In situ* hybridisation for endocardial *gfp* (A,E), myeloid, *spi1b* (C), and myocardial, *myl7* (G), markers in *hand2^uq16ks^* mutants and siblings. Quantification of the expression is shown in B, D, F and H. Dorsal views are shown with anterior to the top in all images. Scale bars represent 100 µm. P values are indicated in graphs.

**Figure 3 supplement 4.**
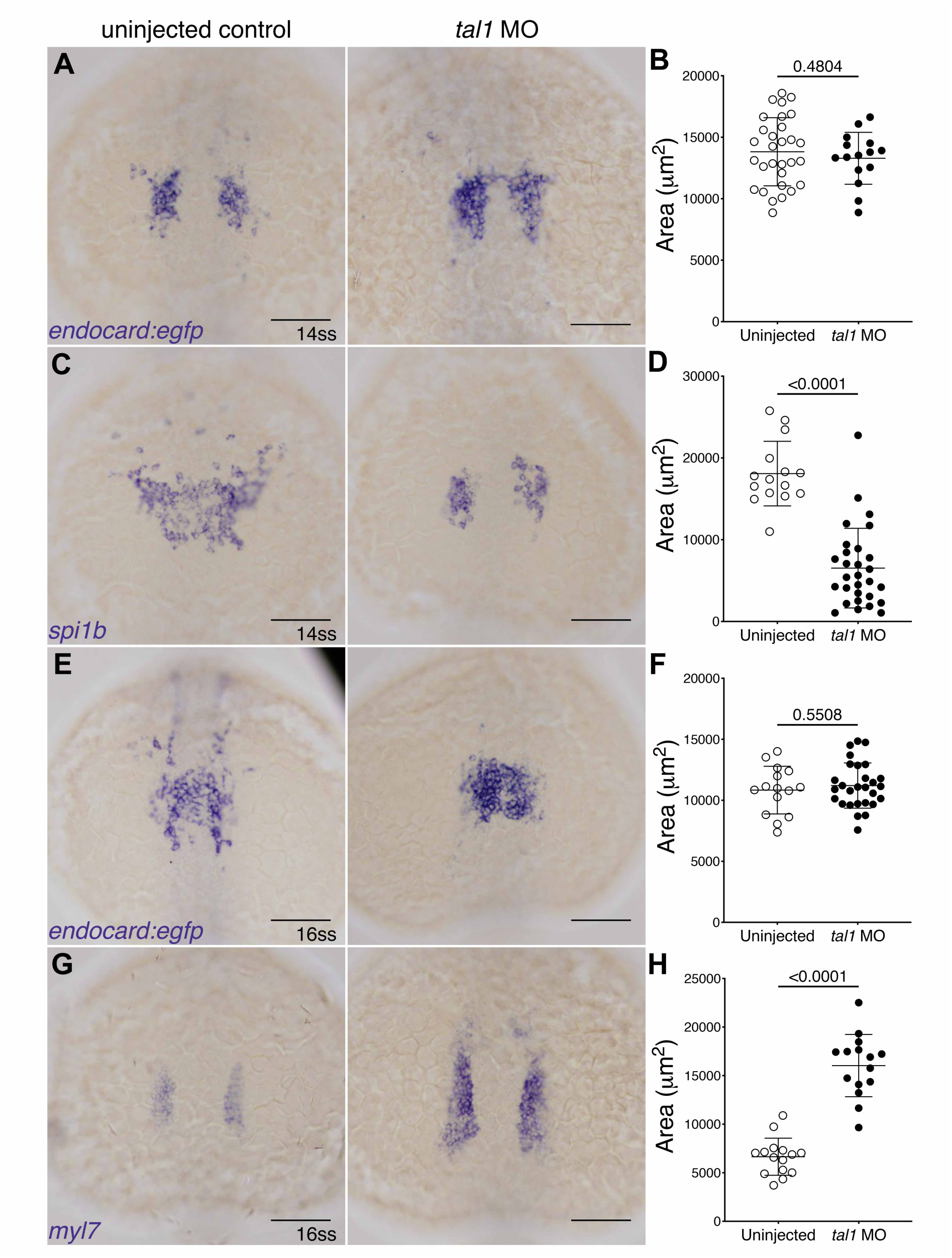
*In situ* hybridisation for endocardial *gfp* (A,E), myeloid, *spi1b* (C), and myocardial, *myl7* (G), markers in *tal1* morphants injected with 0.8 pmol MO and uninjected control (UIC) embryos. Quantification of the expression is shown in B, D, F and H. Dorsal views are shown with anterior to the top in all images. Scale bars represent 100 µm. P values are indicated in graphs.

**Figure 3 supplement 5.**
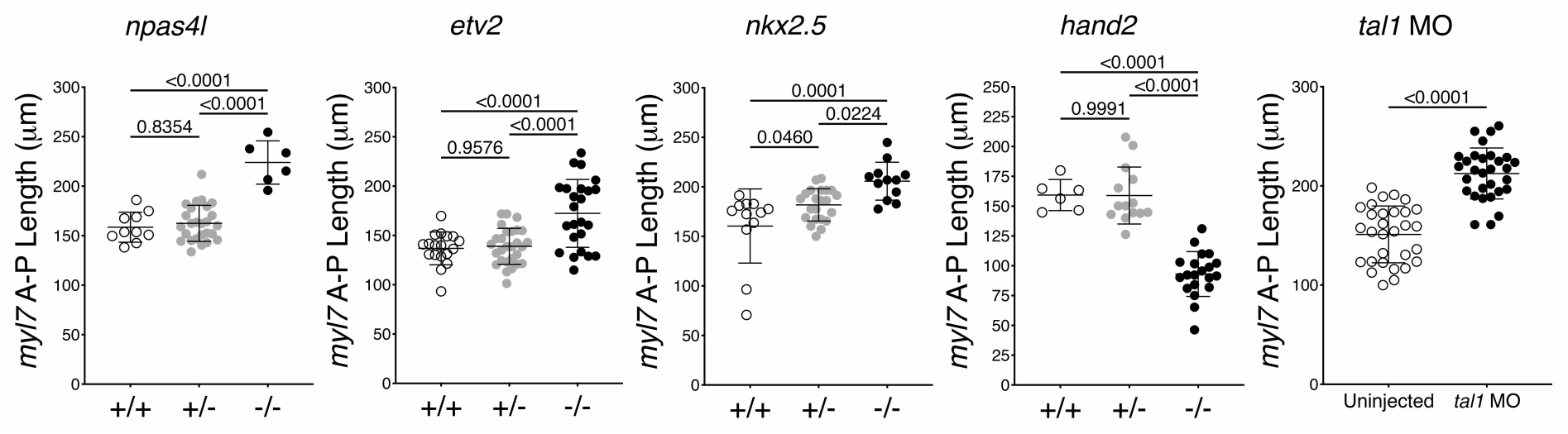
The length of the myocardial expression domains, as defined by *myl7* expression, were measured along the anterior-posterior axis in *npas4l^uq14ks^* (A), *etv2^uq13ks^* (B), *nkx2.5^uq15ks^* (C), and *hand2^uq16ks^*mutants (D) as well as in *tal1* (F) morphants. P values are indicated in graphs.

**Figure 5 supplement 1.**
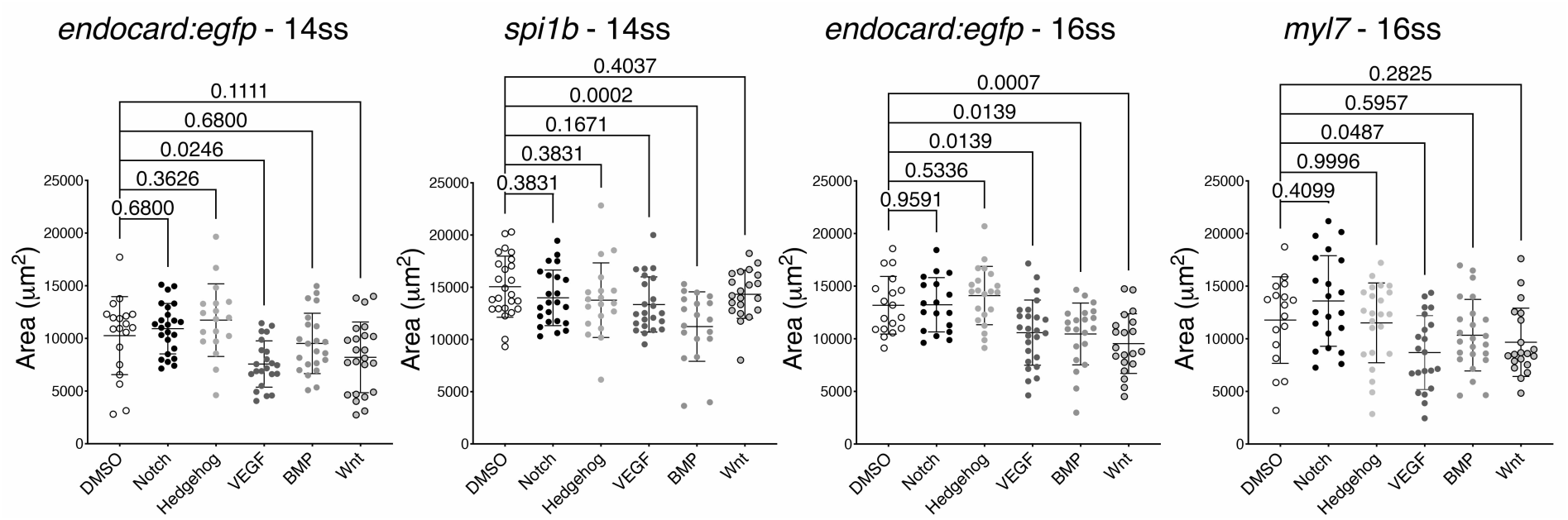
The effect of inhibiting a number of signalling pathways on the formation of endocardial, myeloid and myocardial expression domains as assessed by *gfp*, *spi1b* and *myl7* expression. Drugs were added at the 5ss and embryos incubated until the 14- or 16-ss. Quantification of expression of relevant markers was examined by *in situ* hybridisation staining and surface area measurements as previously described. P values are indicated in graphs.

**Figure 6 supplement 1.**
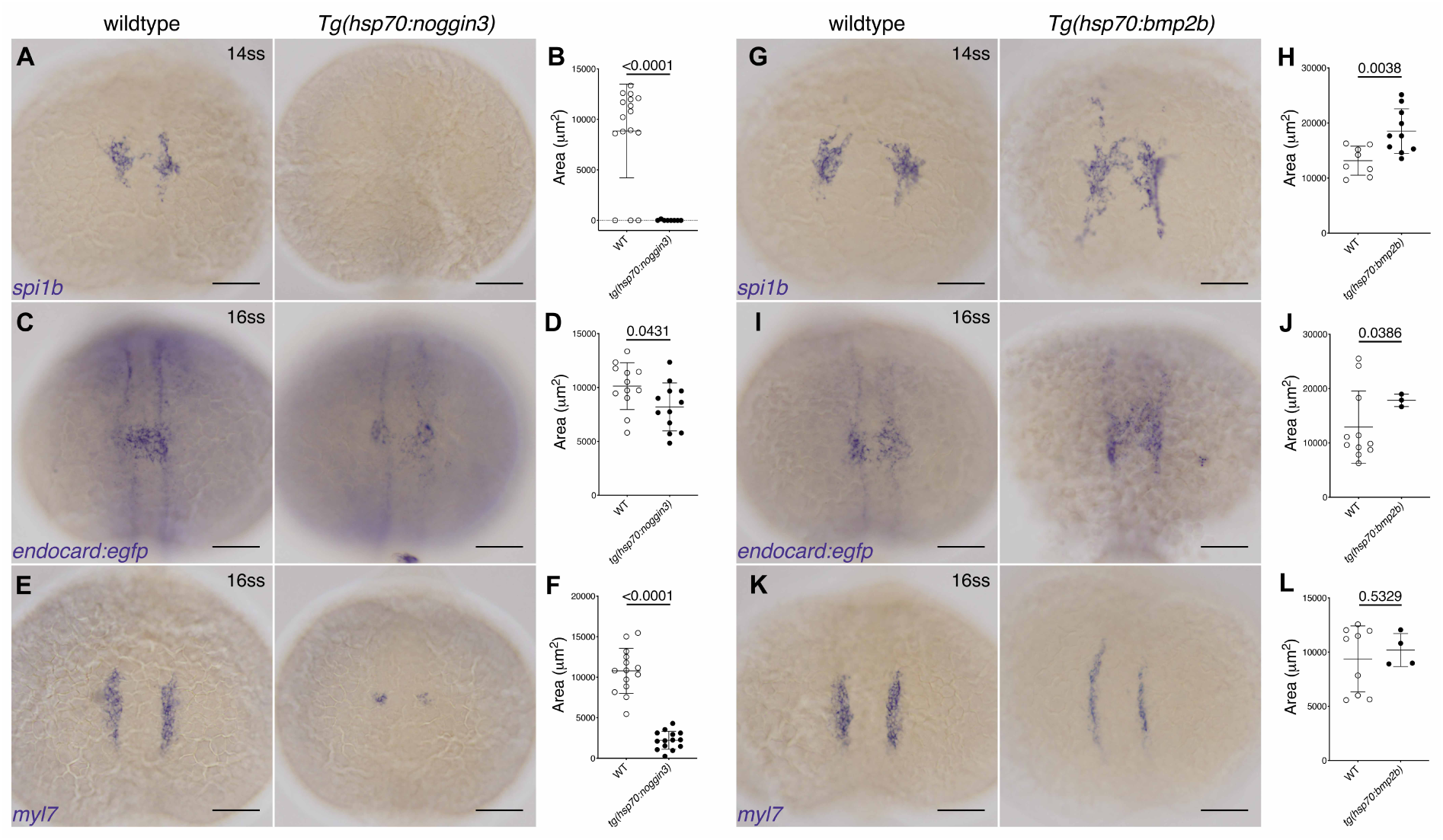
*In situ* hybridisation for myeloid, *spi1b* (A,G), endocardial, *gfp* (C,I), and myocardial, *myl7* (E,K), markers in embryos from crosses of the *Gt(endocard:egfp)* line to either the *Tg(hsp70:noggin3)* or *Tg(hsp70:bmp2b)* lines. Quantification of the expression is shown in B, D, F, H, J and L. Dorsal views are shown with anterior to the top in all images. Scale bars represent 100 µm. P values are indicated in graphs.

